# Network Based Identification of Holistic Drug Target for Parkinson Disease and Deep Learning assisted Drug Repurposing

**DOI:** 10.1101/2022.11.18.515243

**Authors:** Ahsan Raza, Muhammad Muddassar

## Abstract

Parkinson is a neurodegenerative disorder of the nervous system involved with disrupting the motor activity of the body. The current pathogenesis of the disorder is incomplete resulting in widespread use of exogenous medical treatments targeting the dopamine quantity, posing a major challenge in appropriate drug development. The plethora of high throughput techniques in the last decade has yielded a vast amount of Omics dataset with an opportunity of providing a holistic overview of the disease workings and dynamics. We integrated the Parkinson disease Omics datasets using network-based integration strategies to build Parkinson disease network. The most impactful and resilient node of the network was selected as a drug target. Deep learning based virtual screening estimator was built from physicochemical properties of different compounds having variable affinity to target binding. Virtual screening of FDA approved drugs repurposed 19 drugs with 25% of them falling under insomnia treatment; the most prevalent sleep disorder in Parkinson patients. Source Code of the project is available at https://github.com/aysanraza/pd_repurposing_protocol

## Introduction

Parkinson is a complex neurodegenerative disorder of the nervous system that is involved in disrupting the motor and non-motor activity of the human body (Poewe et al., 2017). It was first discovered, reported, and recognized as a medical condition in a monograph of 1817 titled “An Essay on the Shaking Palsy”, by an English surgeon James Parkinson (Toodayan, 2018). It is mostly associated with motor dysfunction of the human body including tremors, stiffness in limbs, slowness in movement, disturbed coordinate balance; non-motor symptoms include depression, hallucinosis, mood disorders, urinary problems, and REM sleep behavior disorder (Poewe et al., 2017). Epidemiology shows it to be rare before the age of 50 but increases 5-10 fold with the 6th to 9th decade of normal human lifespan (Lee & Gilbert, 2016). It is a complex disorder with neuropathology associated with the loss of pigmented area in Substantia nigra pars compacta and Locus coeruleus as well as the aggregation of Lewy-body pathology (Dickson, 2018). Current medical treatments are mostly associated with the exogenic drug administration to balance the dopamine quantity in the affected regions, targeted drug treatments are in their infancy due to an incomplete molecular level pathogenesis (T. K. Lee & Yankee, 2022). New research encompassing OMICS technologies is shedding light over different molecular components of a living cell and disease biology (Hasin et al., 2017). Network Medicine and its methods are in use to better integrate the dataset coming from OMICS technologies to have a more broader prospect of dynamic cellular working and identification of probable drug targets (Dimitrakopoulos et al., 2018). Machine learning based Drug repurposing strategies are in use to identify faster and accurate new drug indications (Tanoli et al., 2021).

Xing & Gardner (2006) developed the first network science based MNI algorithm that takes in gene expression network dataset to infer appropriate drug targets. Sridhar et al. (2006) and Song et al. (2009) utilized the metabolic network dataset to design the respected algorithms that utilize linear approach of backtracking metabolic network until a suboptimal point of desired result is achieved for identification of better performing fast drug target identifications. Kushwaha & Shakya (2010) utilized two fold PPI network analysis to screen impactful and pathologically relevant proteins to screen 18 potential drug targets in Mycobacterium tuberculosis. As drug target identification through network methodologies were in their infancy Ashburn & Thor (2004) started drug repurposing. The advent of drug repurposing was mostly backed by computational analysis including the all important molecular docking studies. The utilization of ML in the domain started with the work of Kinnings et al. (2011) to optimize the docking scores by using ML capabilities. On the other side the development of network based target identification strategies take a sharp turn with the study of Bánky et al. (2013) to utilize Google’s page rank algorithm to better represent the metabolic dataset for appropriate drug target identification. Peng & Schork (2014) follows the further refinement, sees integrity and connectivity of biological networks as an important measure to develop a method that can apply careful centrality measures to screen cancer therapeutic targets. Biological processes are not only interconnected but are dynamic in nature hence Wu et al. (2015) set out to utilize network controllability methods to capture dynamic nature of the biological networks for better screening of potential drug targets. Patrick et al. (2019) make use of NLP word embedding strategy over scientific literature to train drug and disease relationship to screen new drug repositioning in disease states of Immune-mediated diseases. Kuang et al. (2018) utilize electronic medical records to train and predict new drug indications.

Pathogenesis of the Parkinson disease is incomplete, posing a major challenge in the selection of drug target and downstream drug development. Omics sciences on the other hand are providing a vast amount of disease related datasets that, when integrated, can provide a more thorough picture of the internal working of the cell and disorders. This holistic approach towards cellular problems gives us an amazing ability to produce good hypothesis especially in case of a problem with incomplete molecular insights. Network based models are fulfilling the holistic integration of Omics datasets for good hypothesis generation, as shown above network-based strategies to screen or identify drug target molecules for a particular biological problem or disease. The drug target identification can be led forth with utilization of machine learning strategies for faster and accurate drug repurposing, a step forth in targeted drug development for Parkinson treatment.

We will look forward to integrating a disease network model based on its genomics and proteomics datasets. The disease network model will then be subjected to biological annotations for better understanding of the disease and its major modules. Network centrality and robustness analysis will be performed over the network to screen a gene having maximal impact over the network. The maximal effect producing gene will be selected as a drug target and all its known inhibitors alongside a decoy dataset will be extracted and subjected to deep learning based virtual screening estimator building. The trained model will then be further utilized for screening approved drugs for probable candidates drug repurposing.

## Methodology

### Data Extraction and Network Building

GWAS Catalog database was used to extract Genome wide association studies dataset of Parkinson disease on 7^th^ of Jan 2022 (Buniello et al., 2018). STRING database was utilized to screen PPI dataset with the following parameters (Szklarczyk et al., 2020):

- Data Source value of the STRING was only set to *Experimentation* only.
- Confidence level was set to *Low*.
- 1st and 2nd shell values were set to *0* to avoid any Neighbors of the input list.

Cytoscape Software was used to build the disease network containing the SNPs and their respective PPIs; only between them (Shannon et al., 2003).

### Modules Building and Gene Set Enrichment Analysis

The build disease network was then further subjected to the Girvan-Newman Algorithm to extract the major modules of the input network based on the node difference: correspond to the functional effect of the nodes/genes (Girvan & Newman, 2002). Gene Set Enrichment Analysis or GSEA of the build modules of the previous step was conducted through an open source reactome FIViz software package using the datasets of the reactome database. Module’s cutoff was set to 10 and FDR Value to 0.1 to screen major modules for high confidence pathways (G. Wu et al., 2014).

### Centrality and Robustness Analysis

Network-analyzer software plugin of Cytoscape was utilized to implement four major network centrality algorithms including Degree centrality, Betweenness centrality, Closeness centrality and Stress centrality; four network robustness algorithms including Characteristic path length, Average Neighbors, Network density and Network Centralization were utilized to calculate the robustness of the network after manual perturbations (Saito et al., 2012).

### Data Engineering and Feature Extractions

The ChEMBL database was used to extract the known inhibitors of the selected drug target (Gaulton et al., 2016). Strong and Weak IC50 value and decoy dataset (extracted from DUD-E database) was employed to divide the dataset into three major groups (Mysinger et al., 2012). CUI based Mordred software was used to generate 2D/3D descriptors of the build dataset and its respective groups (Moriwaki et al., 2018). Screened molecular descriptors were subjected to Python scripting for cleaning, modification, and rearrangement related reprocessing. SKlearn-based Train test split was used to split the input dataset into training and testing groups. SKlearn-based Standard Scaler was used to standardize the dataset by subtracting the mean and then scaling to unit variance.

### Model Training, Testing and Deployment

SKLearn objects were called to implement major ML algorithms including Multilayer Perceptron, Support Vector Machine, Decision Tree, Random Forest. GridSearchCV algorithm was employed to perform model optimizations for hyperparameter space of the ML algorithms (Pedregosa et al., 2012). The predicted results of the four algorithms were also subjected to four evaluation parameters including Precision, Recall, F1 and Support. Python-based JobLib library was used to pipe the build Deep ML Virtual Screening Estimator for batch deployment.

### Drug Repositioning

*The CHEMBL* database was utilized to extract ATC Level 1 bio-active compounds of the Nervous system on 20^th^ Oct 2022 (Gaulton et al., 2016). *Python* based feature extractions and prepossessing were performed to prepare the dataset to be subjected for predictions. *Bourne-again SHell* command language was used to automate the prediction process for experimental reproducibility and future usage.

**Figure 1:**
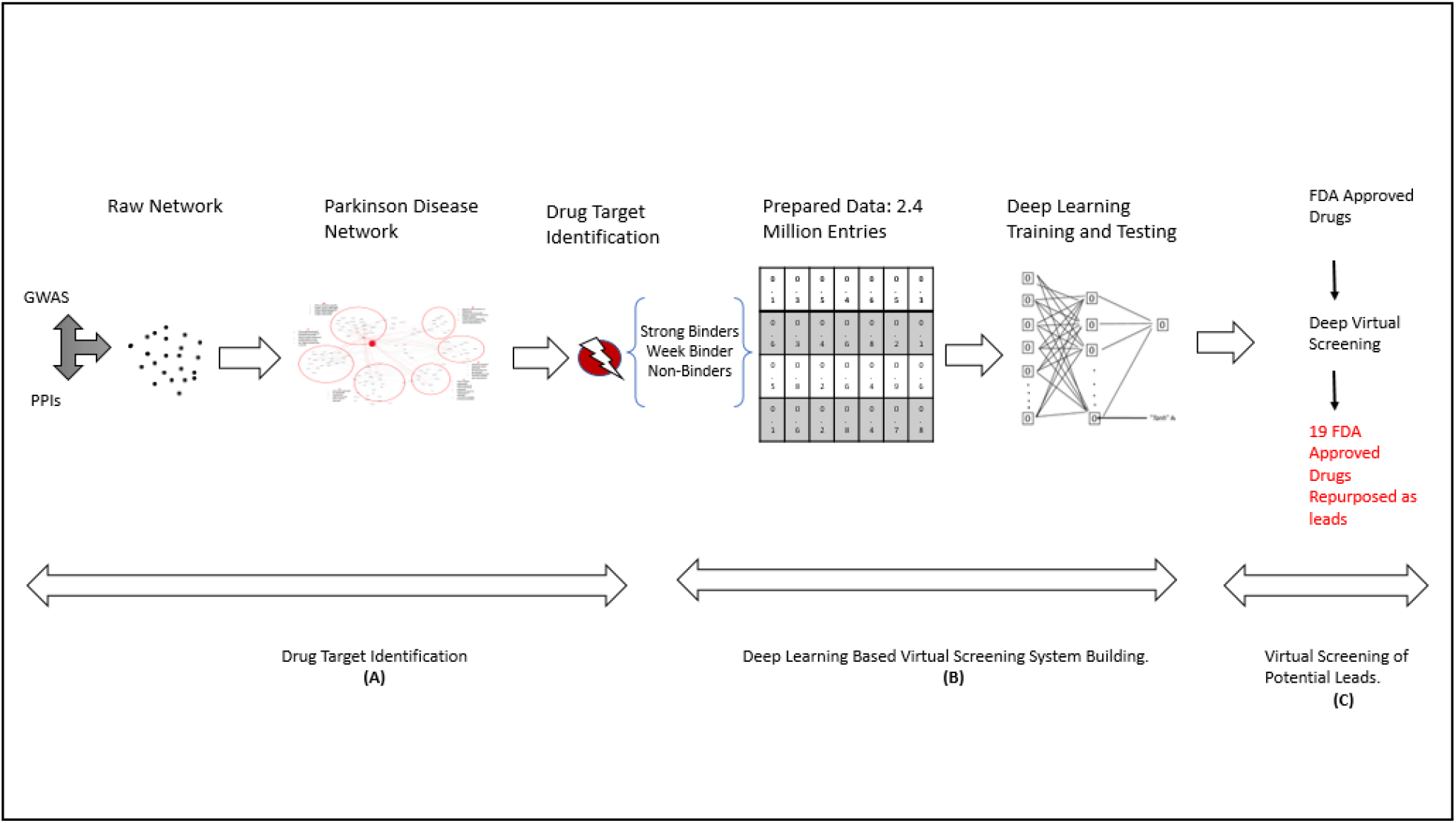
Research Methodology of current study. (A) The dataset coming from genomics and proteomics is integrated towards the building of the Parkinson disease network. Different network communication and connectivity analytics were employed to select the drug target for disrupting the disease network (B) The inhibitors of selected drug targets were divided into three groups and their physicochemical features were used to build a deep learning based virtual screening system. (C) Virtual screening of FDA Approved drugs yields the repurposed drug leads

## Results

### Disease Network

The Genome Wide association dataset of Parkinson Disease contains 54 studies having 505 GWAS associations. The 505 GWAS associations were then subjected to manual cleaning and preprocessing resulting in 270 extracted GWAS associations. A total of 270 SNPs were then subjected to the STRING database and 205 identifiers were mapped in the STRING database against the human protein-protein interaction network. Manual file creation guidelines of the Cytoscape software were used to build a network file based on the genomics and proteomics dataset of the Parkinson disease, imported as a local file into Cytoscape and visualized as a disease network containing 205 nodes corresponding to the filtered SNPs of the Parkinson Disease and 273 edges between them as protein-protein interactions.

### Network Annotation

The build Parkinson disease network containing integrated genomics and proteomics dataset was then subjected to the Girvan-Newman Algorithm to cluster the network into 14 modules, each containing several numbers of nodes corresponding to the genes having similar functional attributes referenced from the reactome database. The disintegrated network with 06 modules was then further subjected to the gene set enrichment analysis of each module through Reactome FIVEz and a total of 420 pathways were screened against the top six major modules of the network. Table 1 enlist the genomic content of six modules and their Gene set enrichment results.

**Table 1.**
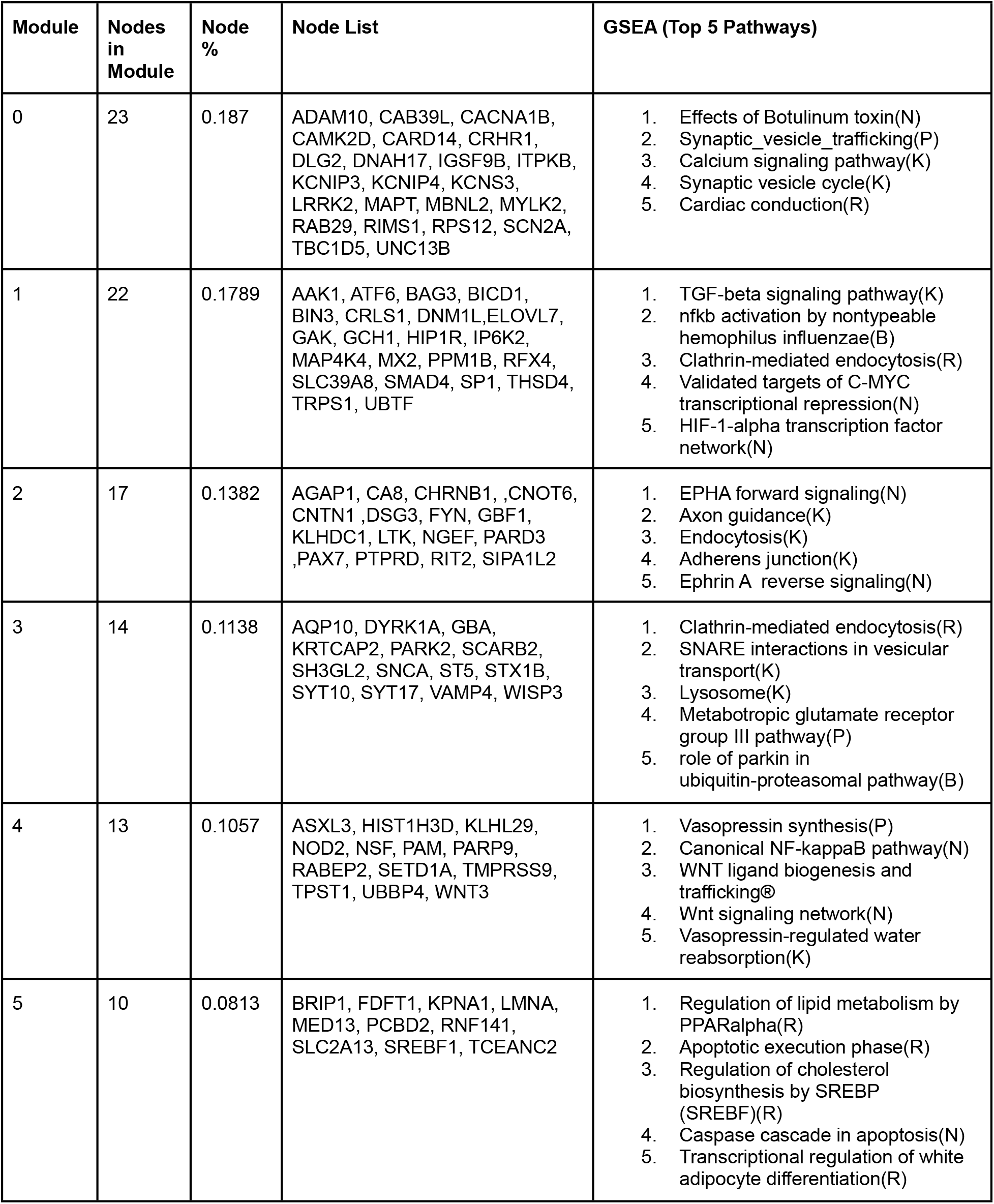
Parkinson disease network modules and their respective nodes and pathways.

### Target Identification

Centrality analysis was performed over the build network to screen the most impactful genes, or the nodes of the network based on their connectivity and communication. Table 2 enlist the top seven candidates of each centrality measure. *SLC2A13, UBBP4, LRRK2, KCNIPA* were predicted as most impact genes in Parkinson disease.

**Table 2.** Centrality analysis of Parkinson disease network.

| No | Betweenness |  | Degree |  | Closeness |  | Stress |  |
| --- | --- | --- | --- | --- | --- | --- | --- | --- |
|  | Gene | Value | Gene | Value | Gene | Value | Gene | Value |
| 1 | FAM134C | 1 | LRRK2 | 33 | SNX29 | 1 | LRRK2 | 14682 |
| 2 | LRRK2 | 0.34 | UBBP4 | 21 | RPS6KL1 | 1 | SLC2A13 | 6966 |
| 3 | SLC2A13 | 0.17 | SLC2A13 | 18 | ZNF608 | 1 | KCNIP4 | 6180 |
| 4 | UBBP4 | 0.14 | DNM1L | 16 | RERE | 1 | UBBP4 | 6026 |
| 5 | KCNIP4 | 0.11 | RIT2 | 15 | FAM134C | 0.8 | GAK | 4210 |
| 6 | HIST1H3D | 0.09 | KCNIP4 | 14 | SV2C | 1 | RIT2 | 3802 |
| 7 | HLA-DRA | 0.07 | GAK | 14 | LRRK2 | 0.509 | HIST1H3D | 3738 |

Network Robustness analysis was performed by manually perturbating a node and seeing an effect it has over the disintegration of the network. Table 3 enlist the values of four robustness measures for selected central nodes. It was found that LRRK2 is dominating in terms of disintegration measures, it has the maximal effect over other nodes as well as modules of the network, have maximal connectivity and communication with highest probability of disintegrating a system.

**Table 3.** Robustness analysis of Parkinson disease network.

| No | Measure | Normal | LRRK2 | SLC2A13 | UBBP4 | KCNIP4 |
| --- | --- | --- | --- | --- | --- | --- |
| 1 | Network Centralization | 0.238 | 0.135 | 0.234 | 0.234 | 0.233 |
| 2 | Characteristic path length | 3.280 | 3.487 | 3.380 | 3.340 | 3.176 |
| 3 | Avg. Number of Neighbors. | 4.439 | 3.934 | 4.180 | 4.131 | 4.246 |
| 4 | Network Density | 0.036 | 0.033 | 0.035 | 0.034 | 0.035 |

### Data Engineering and Feature Extractions

ChEMBL database searched on 13^th^ of March 2022, against the LRRK2 gene, to extract the 1999 inhibitors of the LRRK2 based on IC50 value. Decoy Database was searched on 15^th^ of March 2022, to extract the General Kinase inhibitors as non-inhibitors. Three groups were made based on these dataset as follows:

- Strong Inhibitors: 510 compounds with IC50 value <18nm.
- Week Inhibitors: 510 Compounds with IC50 value >200 nm
- Decoy: 510 randomly selected compounds from decoy dataset.

A total of 1530 compounds were selected with 510 in each group, to lead forward and generate 1614 2D/3D Molecular descriptors from CUI based Mordred tool: making an array containing ~2.4 Million entries. Further data preprocessing led us with an array containing 530 entries and 1376 molecular descriptors to be used by ML models. The dataset was split into training and testing groups with each having 70% and 30% of the data, respectively. Standard Scaler was used to standardize the dataset by subtracting the mean and then scaling to unit variance.

### Deep Learning: Model Building

Dataset was randomly selected for training and testing purposes, 10 times for each algorithm. It was found that the accuracy score of SVM, Random Forest and Decision tree was ~77%, 75%, ~74% respectively. The accuracy score of the deep learning based MLP was ~80%; which is almost 3-4% more accurate than the traditional ML algorithms. Table 4 enlist the accuracy score of the algorithms.

**Table 4.** Accuracy results of random experimentations across build models.

| Accuracy Test No. | MLP | SVM | Random Forest | Decision Tree |
| --- | --- | --- | --- | --- |
| 1 | 79 | 74 | 68 | 73 |
| 2 | 77 | 78 | 74 | 77 |
| 3 | 81 | 74 | 73 | 73 |
| 4 | 79 | 77 | 78 | 69 |
| 5 | 82 | 79 | 75 | 75 |
| 6 | 79 | 82 | 77 | 74 |
| 7 | 78 | 72 | 76 | 76 |
| 8 | 80 | 80 | 76 | 76 |
| 9 | 79 | 73 | 74 | 73 |
| 10 | 82 | 80 | 75 | 75 |
| **Avg** | **79.6** | **76.9** | **74.6** | **74.1** |

Model evaluation parameters including Precision, Recall, F1 and Support, showed the performance of MLP and SVM were equivalent and better against the predictions of multi labels compared to Decision Tree and Random Forest. Table 5 enlist results of four evaluation parameters.

**Table 5.** Evaluation analysis across the prediction classes for major algorithms.

| Evaluation / Class | MLP | SVM | Decision Tree | Random Forest |
| --- | --- | --- | --- | --- |
| Precision / Strong | 0.64 | 0.64 | 0.66 | 0.65 |
| Precision / Week | 0.67 | 0.67 | 0.65 | 0.68 |
| Precision / Non | 0.99 | 0.99 | 0.95 | 0.90 |
| Recall / Strong | 0.74 | 0.74 | 0.70 | 0.67 |
| Recall / Week | 0.57 | 0.57 | 0.61 | 0.60 |
| Recall / Non | 0.99 | 0.99 | 0.94 | 0.99 |
| F1 / Strong | 0.69 | 0.69 | 0.68 | 0.66 |
| F1 / Week | 0.62 | 0.62 | 0.63 | 0.64 |
| F1 / Non | 0.99 | 0.99 | 0.94 | 0.95 |
| Support / Strong | 161 | 161 | 161 | 161 |
| Support / Week | 148 | 148 | 148 | 148 |
| Support / Non | 149 | 149 | 149 | 149 |

### Drug Repositioning

*The CHEMBL* database was utilized to extract 512 ATC Level 1 bio-active compounds of the Nervous system on 20^th^ Oct 2022. These compounds were subjected to our build LRRK2 Virtual screening system. A total of 62 bio-active compounds were screened as strong inhibitors of LRRK2 with 19 compounds being the approved/withdrawn FDA approved drugs. Table 6 enlist the 19 repurposed FDA drugs and their details.

**Table 6.**
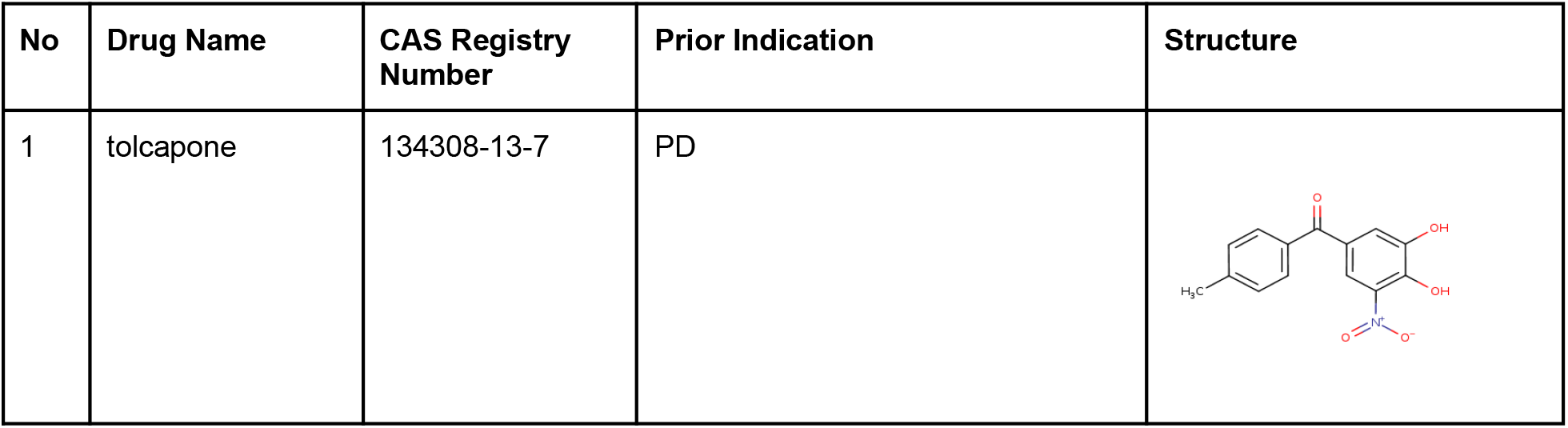
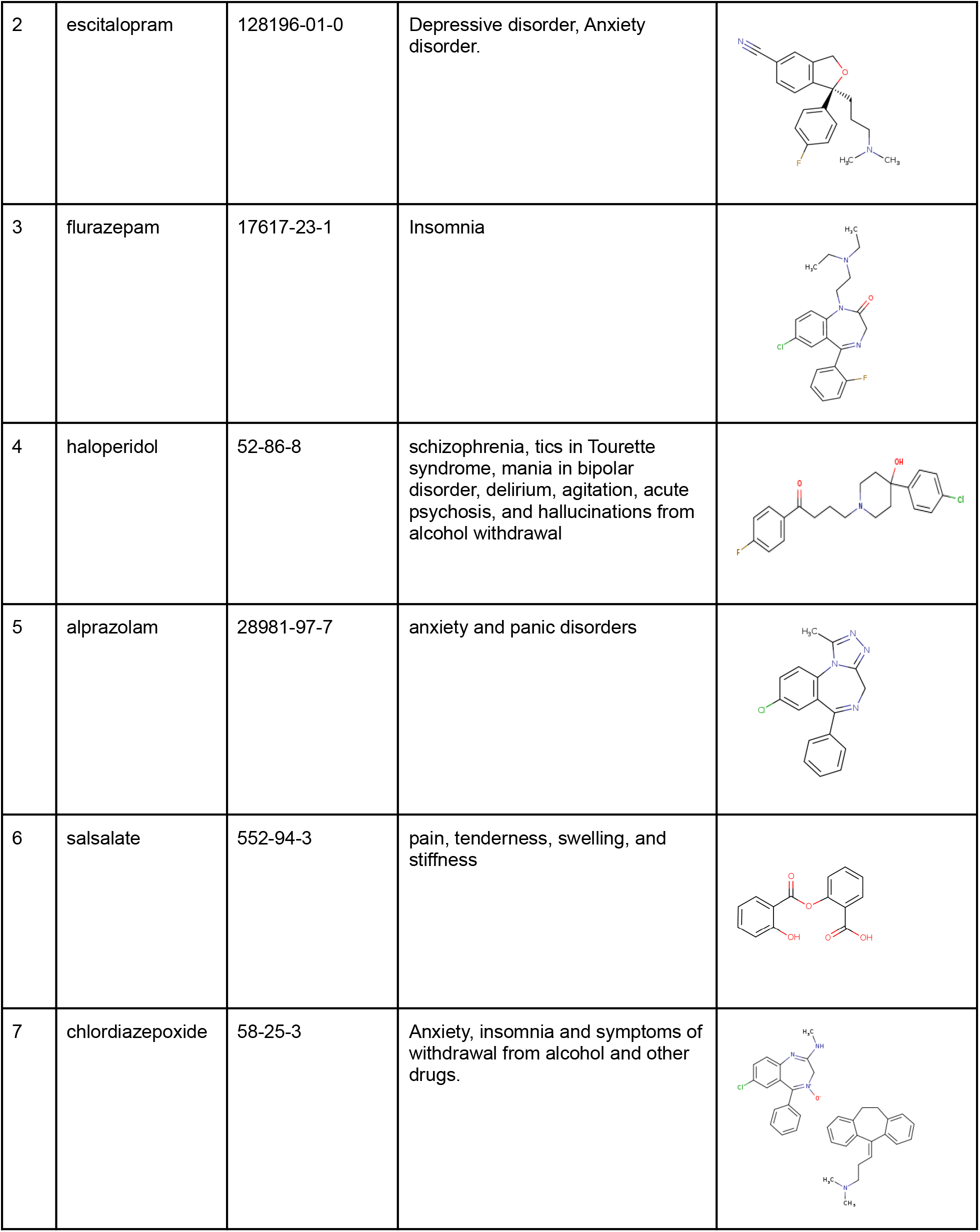
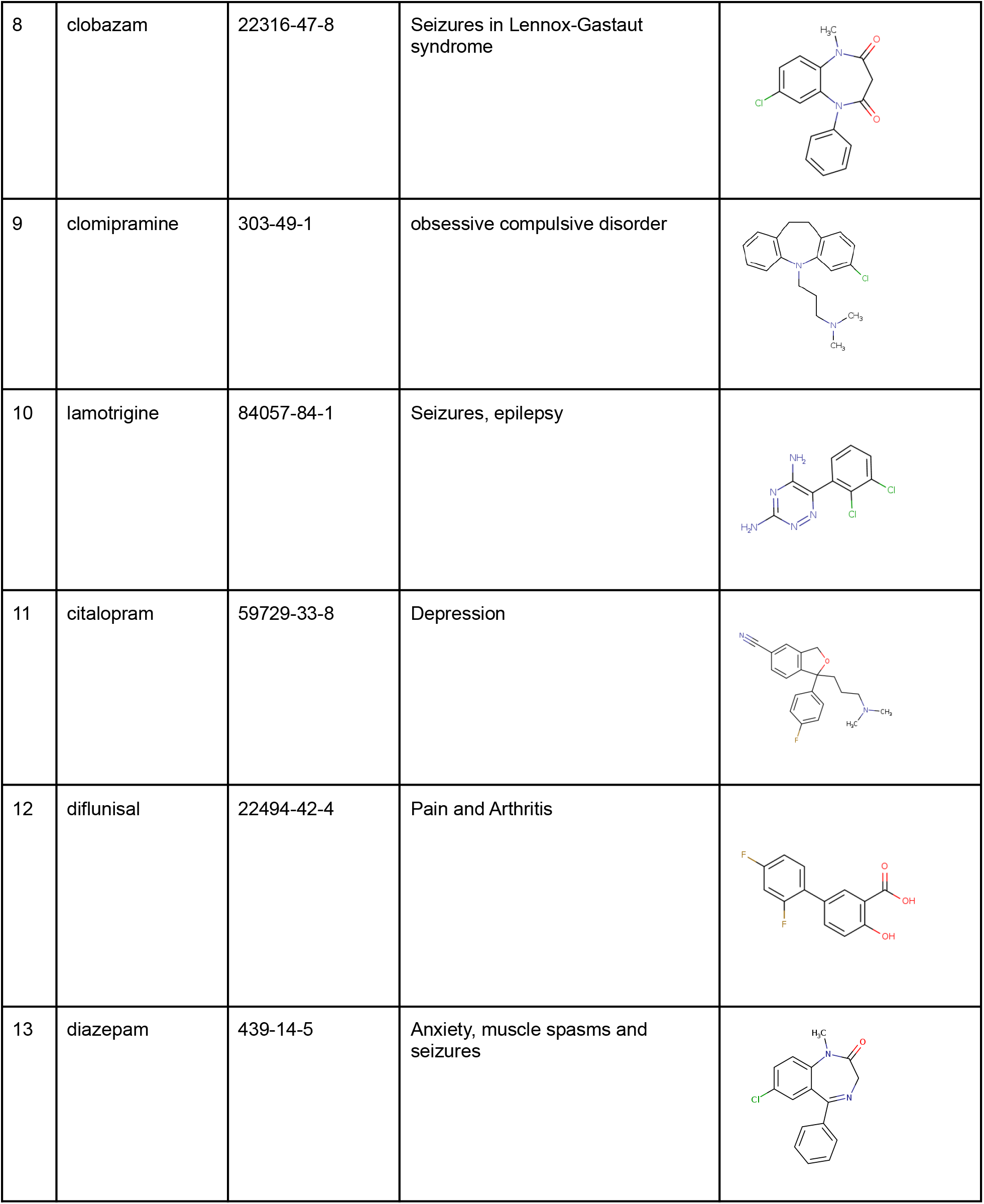
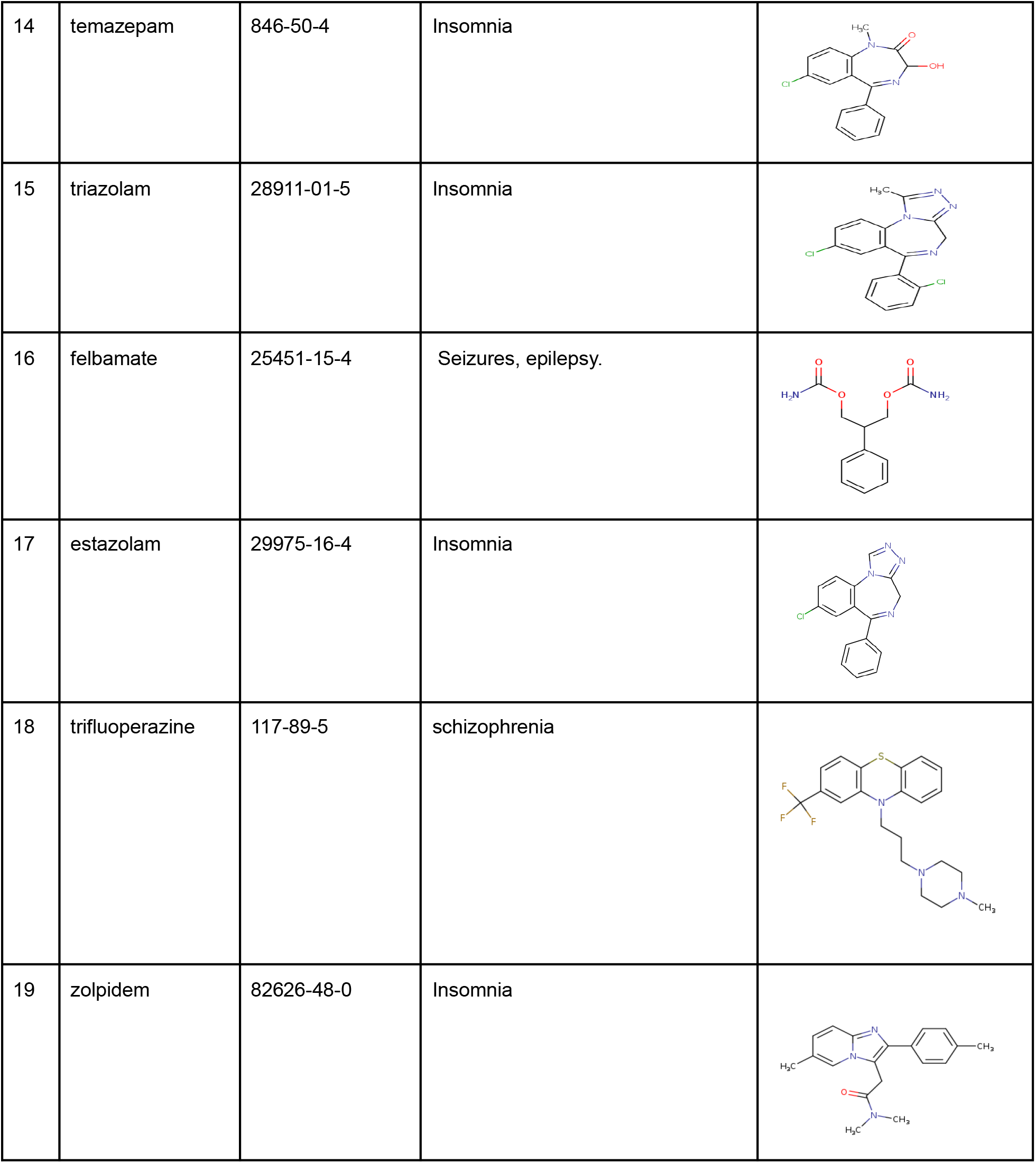
Repurposed drug leads for Parkinson disease.

| No | Drug Name | CAS Registry Number | Prior Indication | Structure |
| --- | --- | --- | --- | --- |
| 1 | tolcapone | 134308-13-7 | PD |  |
| 2 | escitalopram | 128196-01-0 | Depressive disorder, Anxiety disorder. |  |
| 3 | flurazepam | 17617-23-1 | Insomnia |  |
| 4 | haloperidol | 52-86-8 | schizophrenia, tics in Tourette syndrome, mania in bipolar disorder, delirium, agitation, acute psychosis, and hallucinations from alcohol withdrawal |  |
| 5 | alprazolam | 28981-97-7 | anxiety and panic disorders |  |
| 6 | salsalate | 552-94-3 | pain, tenderness, swelling, and stiffness |  |
| 7 | chlordiazepoxide | 58-25-3 | Anxiety, insomnia and symptoms of withdrawal from alcohol and other drugs. |  |
| 8 | clobazam | 22316-47-8 | Seizures in Lennox-Gastaut syndrome |  |
| 9 | clomipramine | 303-49-1 | obsessive compulsive disorder |  |
| 10 | lamotrigine | 84057-84-1 | Seizures, epilepsy |  |
| 11 | citalopram | 59729-33-8 | Depression |  |
| 12 | diflunisal | 22494-42-4 | Pain and Arthritis |  |
| 13 | diazepam | 439-14-5 | Anxiety, muscle spasms and seizures |  |
| 14 | temazepam | 846-50-4 | Insomnia |  |
| 15 | triazolam | 28911-01-5 | Insomnia |  |
| 16 | felbamate | 25451-15-4 | Seizures, epilepsy. |  |
| 17 | estazolam | 29975-16-4 | Insomnia |  |
| 18 | trifluoperazine | 117-89-5 | schizophrenia |  |
| 19 | zolpidem | 82626-48-0 | Insomnia |  |

## Discussion

### Parkinson Disease Network

Parkinson is a neurodegenerative disorder of the nervous system involved in disrupting the motor activity of the human body. There is a limitation of conventional drug therapeutic interventions of Parkinson, majorly due to incomplete molecular pathogenesis of the disease (Poewe et al., 2017). Omics datasets from the last decade and invention of its integration strategies provide means to understand folded realities of biological workings (Subramanian et al., 2020). We integrated the Genomics and Proteomics dataset of Parkinson disease into a disease Network for further network base drug target identifications but before leading forth, we disintegrated our network into different identical modules and performed enrichment analysis over each module to confirm the validity of the build network as a disease network. The enrichment of Calcium, Cellular signaling, endocytosis, Lysosomal dysfunction, Signal Transduction, cellular response and Lipid Metabolism related pathways in our result and their respective association with Parkinson’s incident, progression and development is well established (Surmeier et al., 2017; Fujita et al., 2013; Vidyadhara et al., 2019; Hoang, 2014; Baekelandt et al., 2020). The build disease network shows striking similarities with the disease annotations hence confidently acclaimed as a Parkinson disease network. Figure 2 shows the complete Parkinson Disease network and its associated modules and their enrich pathways.

**Figure 2.**
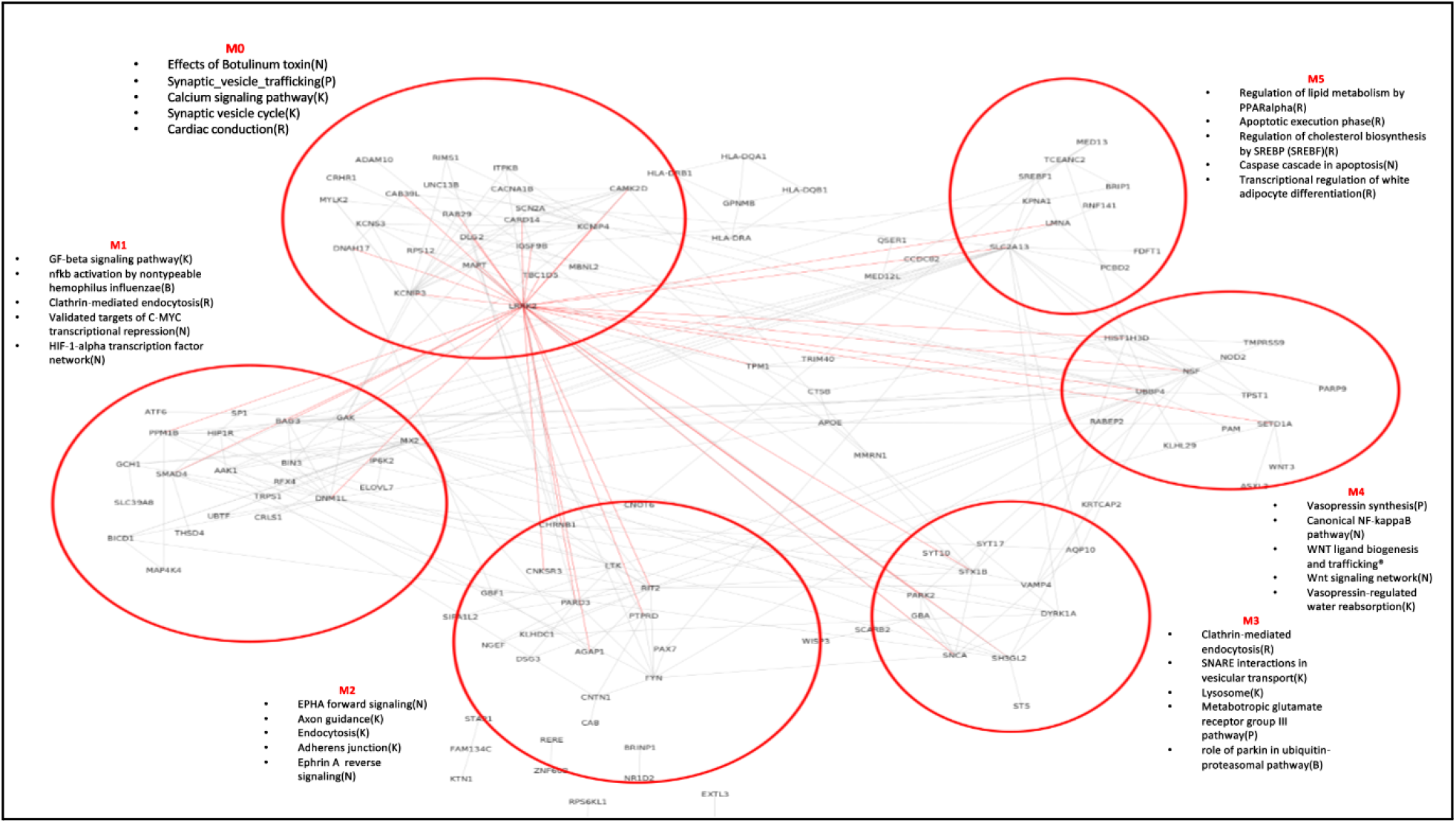
The Complete Parkinson Disease Network. The build disease network and its major 6 modules are enriched with major pathways majorly falling under cell signaling, lipid acid metabolism, degradation systems, calcium homeostasis and apoptosis.

### Drug Target Identification

The build disease network is composed of 204 nodes and 273 edges representing the genomics and proteomics representation of Parkinson disease. Network centrality is a measure of how central a node is in its network and centrality algorithms applied over the disease network shows *SLC2A13, UBBP4, LRRK2, KCNIPA* as the most central and controlling nodes of the network. Figure 3 summarizes the centrality analysis.

**Figure 3.**
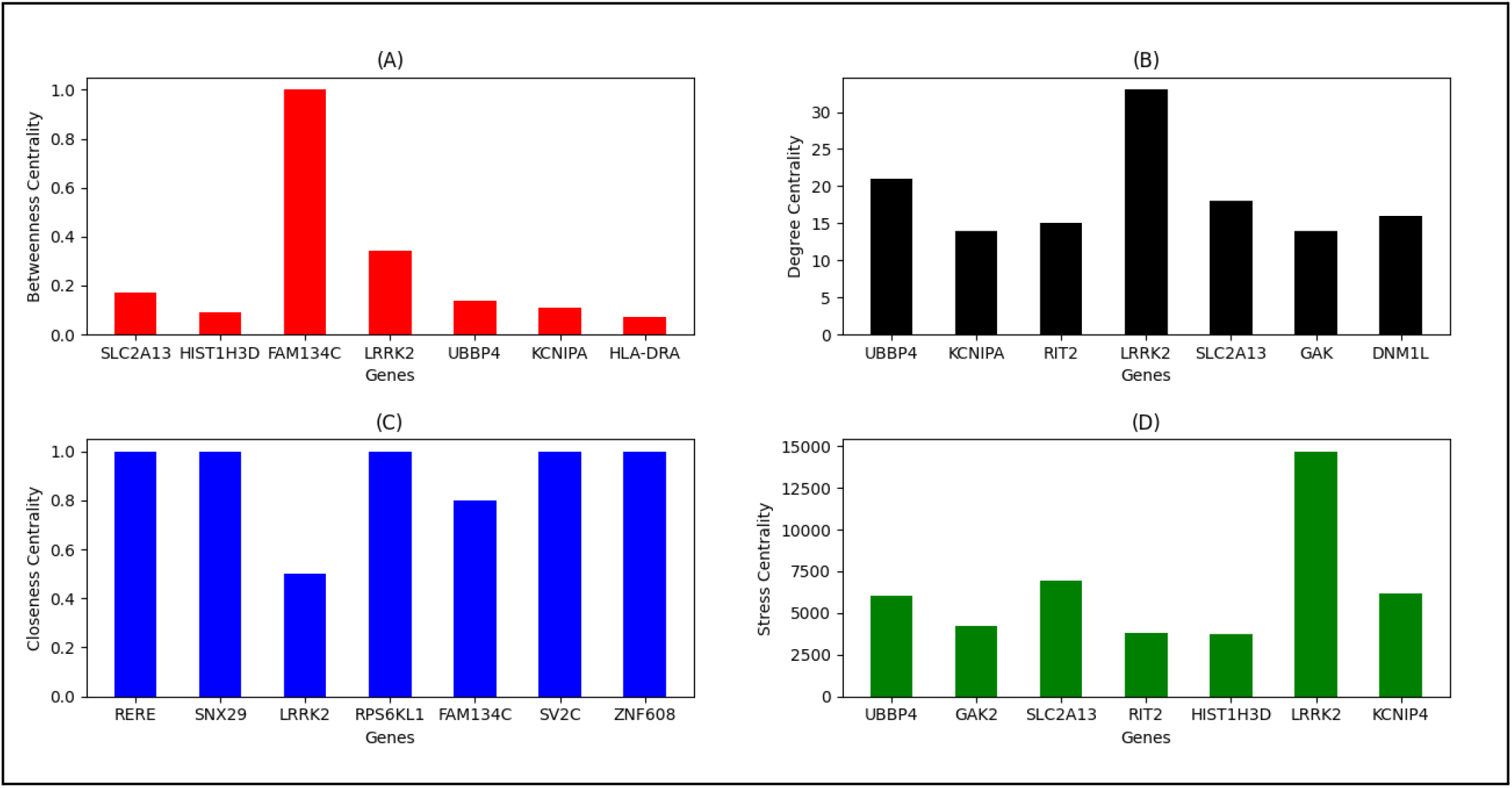
The centrality analysis of the Parkinson disease network. (A) The betweenness centrality analytics showing the top candidate nodes, conducting most of the network’s inter-component communication. (B) The degree of centrality analytics showing the top candidates for most number of connections. (C) The closeness centrality analytics showing the top candidates to absorb the shortest paths of the network. (D) The stress centrality analytics showing the comparative stress absorbing ability of the network communication in top candidates.

In case of disrupting the network, the screening of the most impactful genes of the network in the previous step does not give clear indications of the most consolidating node of the network that resists network disintegration. Hence the most central nodes extracted from the previous step were perturbed from the network and the resulting networks were subjected to four network robustness indicating algorithms. In case of decreasing network density, average number of neighboring nodes, network centralization and increasing the average path of any node to the other, LRRK2 perturbed network shows most deviation compared to the other candidates. It was evident, the perturbation of *LRRK2* from the network yields the most devastating effects over network integration and leads towards disease network disintegration. We hypothesized an overexpressed LRRK2 as the most probable drug target candidate from the above analytics, if inhibited, could cause the most devastating effects over Parkinson disease and its progression. Figure 4 summarizes the network robustness analysis.

**Figure 4.**
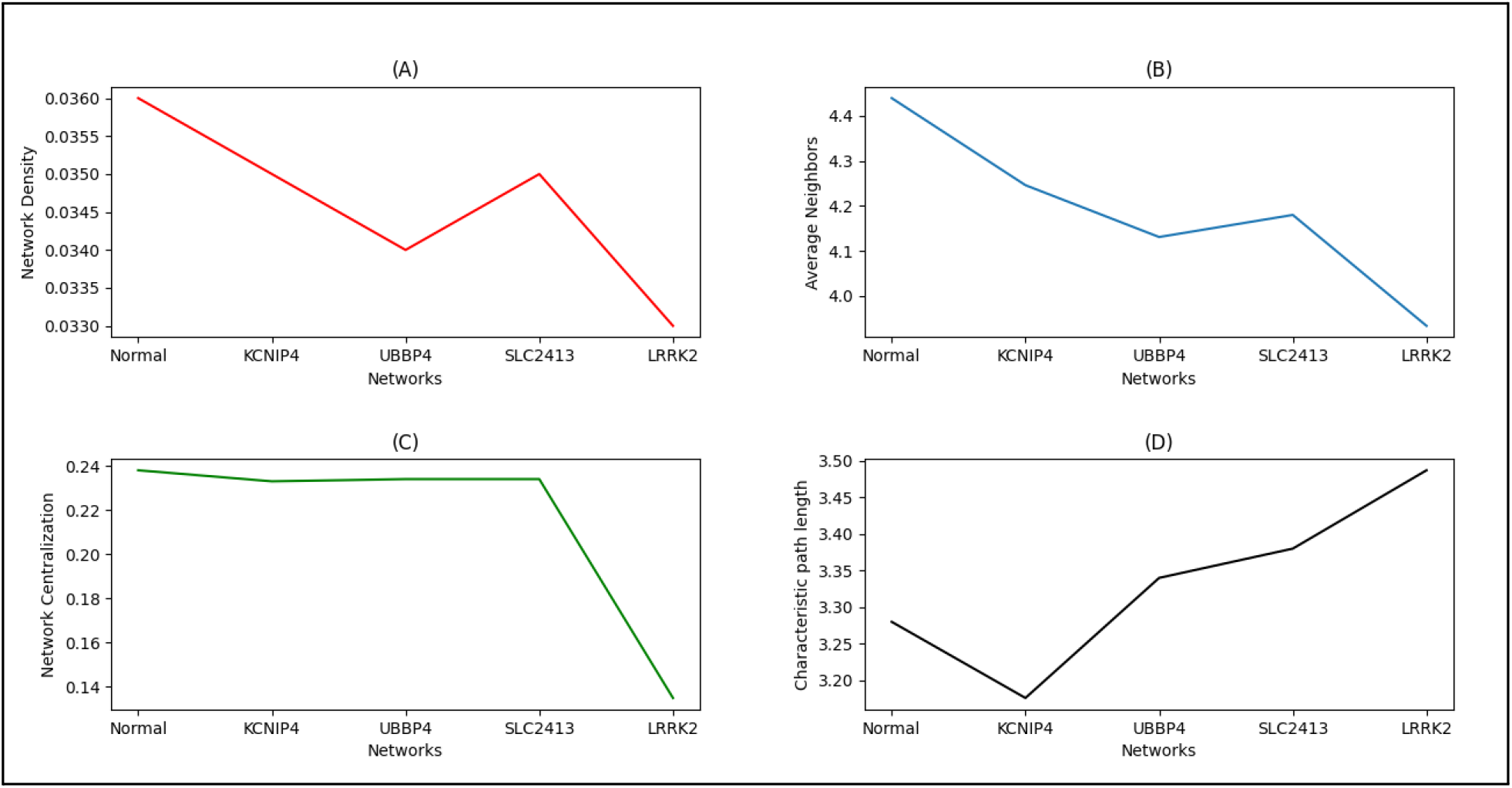
The robustness analysis of Parkinson disease network. (A) The network density analytics showing the LRRK2 perturbation network to have the least network density ratio compared to normal, an indicator of decreased connectivity. (B) The network Average Neighbors analytics showing the LRRK2 perturbation network to have the most decreased neighbors connectivity compared to normal. (C) The network centralization analysis shows a drastic decrease in the centralization of the network compared to a normal disease network, an indicator of disruption in normal disease dynamics (D) The network Characteristic path length analytics showing the LRRK2 perturbation network to have the most decreased communication compared to normal.

### Deep Learning Virtual Screening Estimator

The build virtual screening machine learning model from MLP architecture shows equivalent performance to SVM and better against Random Forest and Decision Tree, when measured across evaluation parameters including Precision, Recall, F1 and Support. The evaluation analysis showed the learning of “non-inhibitor” prediction as most accurate compared to other labels, majorly because of the variation in non-binders dataset. The performance of “strong inhibitors” prediction can be accredited to the dataset limitation of LRRK2 inhibitors. Figure 5 summarizes the evaluation analysis.

**Figure 5.**
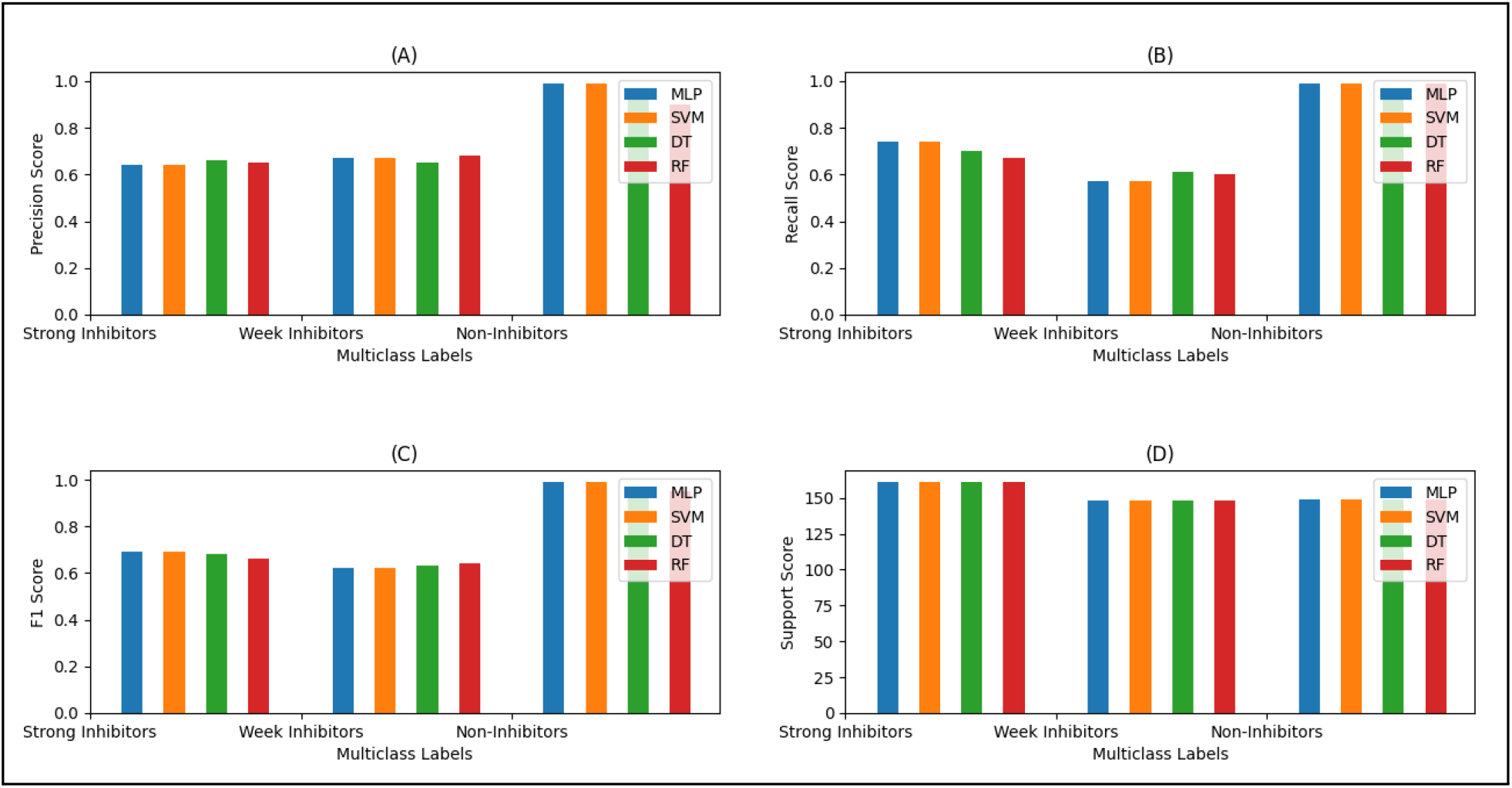
Comparative models evaluation for prediction classes. (A) The precision analytics indicating positive predictions shows the learning of the “non-inhibitor” class as near to best compared to other two classes. The learning of MLP is slightly low against other ML models in case of “strong inhibitors” prediction. (B) The Recall analytics shows the learning of “non-inhibitor’ class as best compared to other two classes. The learning of MLP and SVM is slightly higher against other ML models in case of “strong inhibitors” prediction. (C) The F1 analytics shows the learning of the “non-inhibitor’ class as best compared to other two classes. The learning of MLP and SVM is slightly higher against other ML models in case of “strong inhibitors” prediction. (D) The support analytics shows the learning of all classes as best. The learning of MLP and other ML models in case of “strong inhibitors” prediction is equivalent.

The same MLP model when trained and tested randomly 10 times shows ~80% accuracy compared to ~77% of SVM, ~75% of Random Forest and ~74% of Decision Tree. Figure 6 summaries the average accuracy scores of algorithms. The slightly better performance of MLP architecture can be credited to its ability to capture nonlinear relationships of physicochemical properties of drug compounds without being feature-reduced.

**Figure 6.**
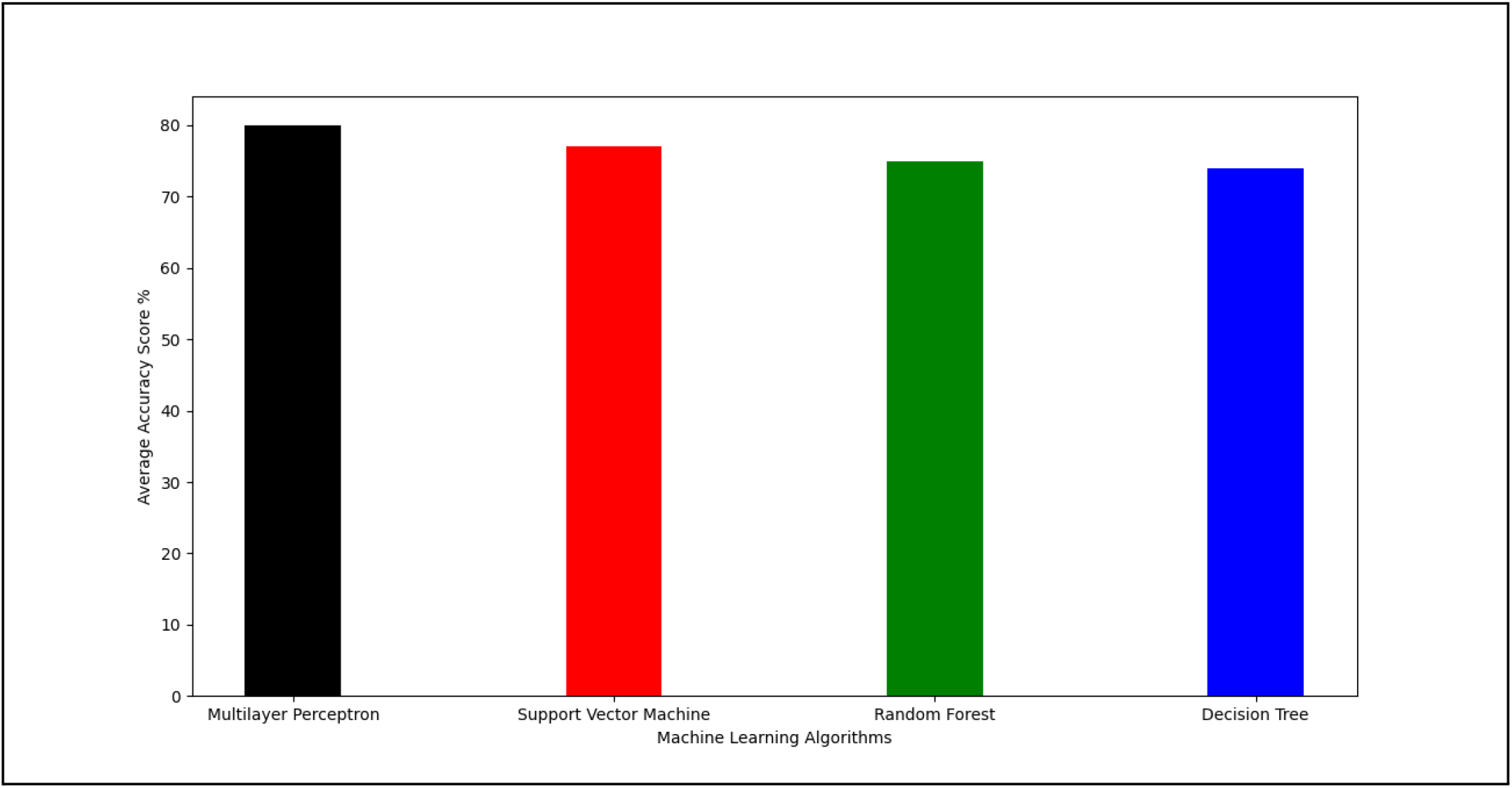
Average Accuracy Analysis. The random accuracy measure of the four models when shuffled, trained and predicted for 10 times shows MLP to perform better at ~80% accuracy compared to SVM, Random Forest and Decision Tree at 77%, 75% and 74% respectively.

We utilized the simple MLPGS class of SkLearn Library to import a simple neural network model. The model was given 1376 molecular descriptors as input in the first layer, with each neuron of the network utilizing the “tanh” activation function to process the neuronal input for reliable output generation. The second layer of 10 neurons were added to further process the input and proceed towards the final and third layer containing three neurons falling under three major choices in our case: non-binders, weak binders and strong binders. The powerful “adam” optimizer was added to backpropagate the architecture and reset the network weights for output tuning. The initial learning rate for weights was set at “0.001” with learning rate as “adaptive”. The maximum number of iteration for re-tuning the output was set up to 2000; to keep the utilization of computational resources and reliable learning in consideration. Figure 7 shows the architecture of our build Multilayer Perceptron Model.

**Figure 7.**
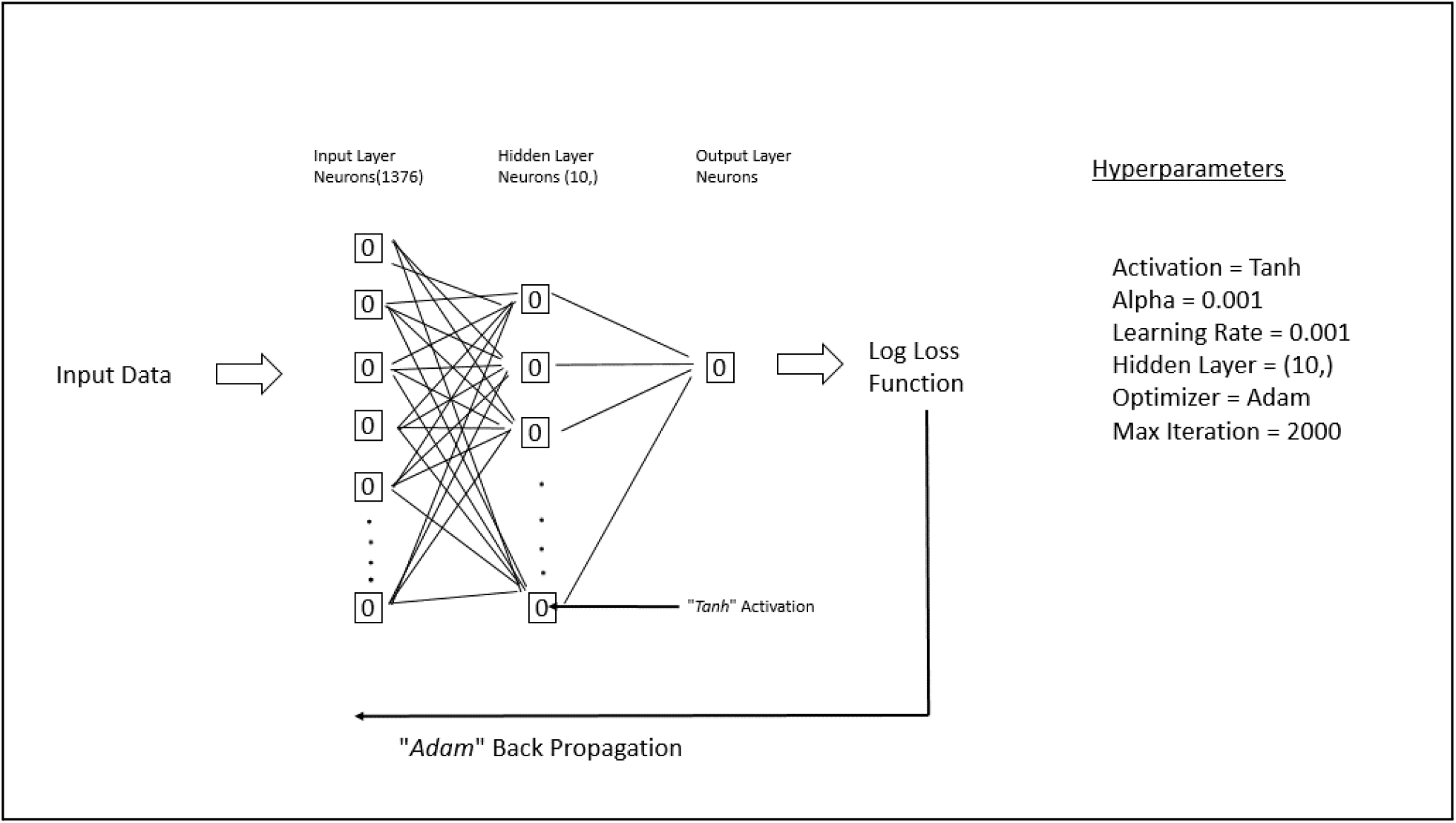
The utilized Neural Network Architecture of Multi Layer Perceptron. The build neural network contained input and output layers, and a hidden layer containing 10 neurons. *Tanh* activation function is utilized with *alpha* and *learning rate* set to 0.001. The *log loss* calculation is followed with *Adam* optimization for back propagation and resetting the weights. The architecture is limited to iterating the incoming dataset for no more than 2000 times.

### Drug Repositioning

The deep learning based virtual screening system we built yielded 19 FDA drugs as repurposed candidates that have the potential to inhibit LRRK2 as targeted drug leads. The 25% of the repurposed leads fall under the insomnia treatment therapeutics. As insomnia is the most prevalent sleep related diagnosed problem in Parkinson patients, we believe in the utilization of further trails as safe and better for this 25% of drug groups containing “*flurazepam*”, “*temazepam*”, “*triazolam*”, “*estazolam*”, and “*zolpidem*”. Figure 8 shows the summary of screened drug leads and their respective indications.

**Figure 8.**
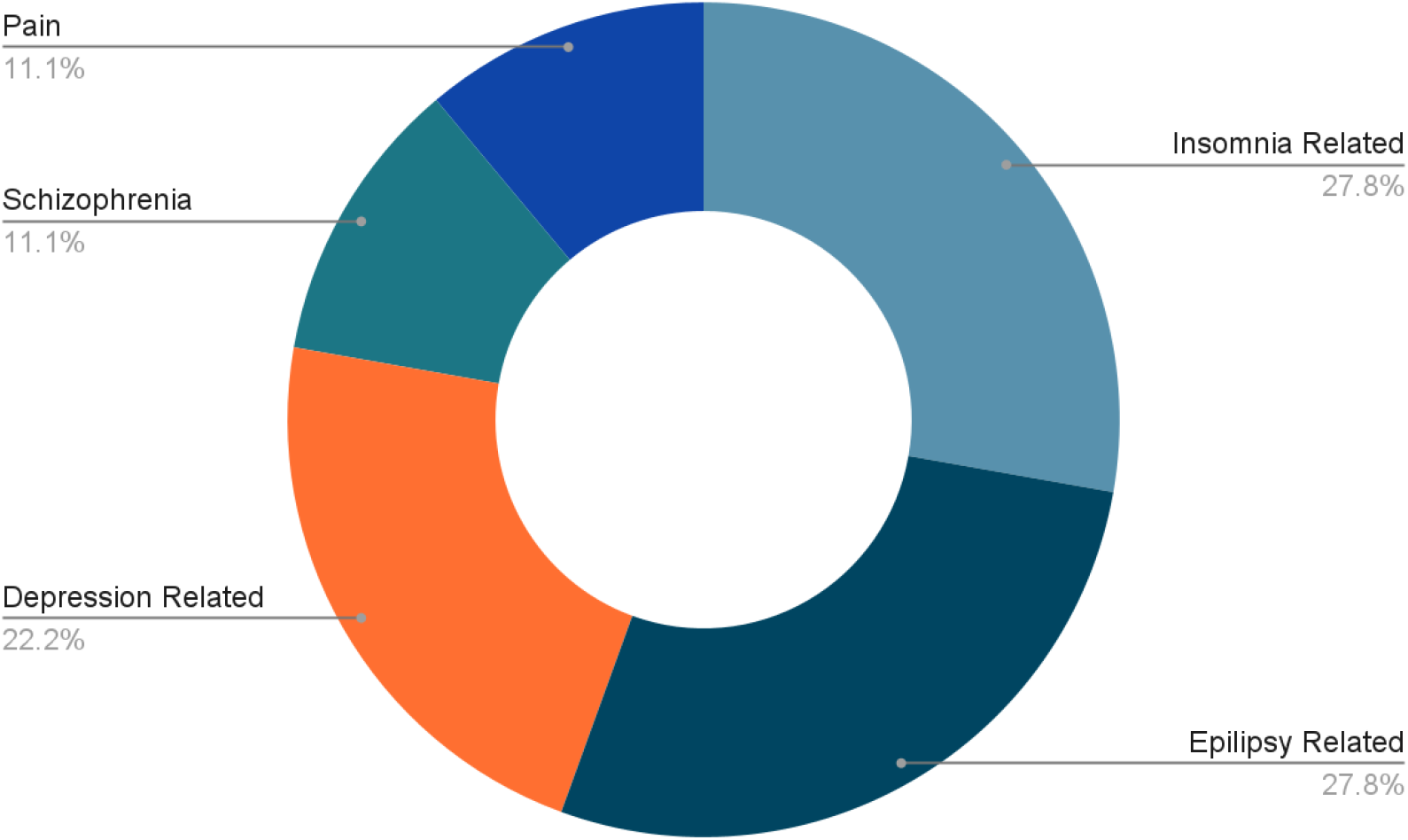
The summary of indication classes for repurposed drugs. Insomnia and Epilepsy related indications are prevalent with 55% of the group. Depression, Schizophrenia and pain related indications comprise the remaining 45% of the screen drugs.

### Limitations and Prospects

We want to report few of the limitations and future prospects of this study:

1. We utilize the Genomics and Proteomics dataset for disease network building but that alone does not represent the whole biological fingerprint of a cell and utilization of other OMICS datasets can provide more interesting results.
2. We utilize the MLP based deep learning architecture for virtual screening estimator building but the utilization of other deep architecture like CNN or RNN might give better accuracy.
3. The current study tries to find the holistic drug target to leadforth with targeted drug therapeutics but lacks the research work on the potential genomics side effects associated with targeting LRRK2 based therapy.
4. There is a need for further studies regarding the genomic interrelations of the prior-indications of proposed drug leads and Parkinson disease.

### Conclusion

Parkinson is a neurodegenerative disorder of the nervous system with incomplete molecular pathogenesis; posing a major challenge in targeted drug therapy. We integrated semi-OMICS disease datasets through network strategies to build the disease network and utilize network based centrality and robustness analytics to hypothesize LRRK2 as a holistic drug target. Deep learning based virtual screening estimator was built from physicochemical properties of different compounds having IC50 based variable affinity to LRRK2 binding. The build virtual screening system was given with the ATC level 1 compounds of the nervous system to screen 19 FDA drugs as potential repurposed leads for targeted drug therapy. The current study has dataset and accuracy based limitations but is a substantial systematic approach towards targeted drug development for a disease with challenges of incomplete molecular pathogenesis.

## References

Ashburn, T. T., & Thor, K. B. (2004). Drug repositioning: Identifying and developing new uses for existing drugs. Nature Reviews Drug Discovery, 3(8), 673–683. https://doi.org/10.1038/nrd1468

Baekelandt, V., Lobbestael, E., Xicoy, H., & Martens, G. J. M. (2020). Editorial: The role of lipids in the pathogenesis of parkinson’s disease. Frontiers in Neuroscience, 14. https://doi.org/10.3389/fnins.2020.00250

Bánky, D., Iván, G., & Grolmusz, V. (2013). Equal opportunity for low-degree network nodes: A pagerank-based method for protein target identification in metabolic graphs. PLoS ONE, 8(1), e54204. https://doi.org/10.1371/journal.pone.0054204

Buniello, A., MacArthur, J. A. L., Cerezo, M., Harris, L. W., Hayhurst, J., Malangone, C., McMahon, A., Morales, J., Mountjoy, E., Sollis, E., Suveges, D., Vrousgou, O., Whetzel, P. L., Amode, R., Guillen, J. A., Riat, H. S., Trevanion, S. J., Hall, P., Junkins, H., … Parkinson, H. (2018). The NHGRI-EBI GWAS Catalog of published genome-wide association studies, targeted arrays and summary statistics 2019. Nucleic Acids Research, 47(D1), D1005–D1012. https://doi.org/10.1093/nar/gky1120

Dickson, D. W. (2018). Neuropathology of Parkinson disease. Parkinsonism &amp; Related Disorders, 46, S30–S33. https://doi.org/10.1016/j.parkreldis.2017.07.033

Dimitrakopoulos, C., Hindupur, S. K., Häfliger, L., Behr, J., Montazeri, H., Hall, M. N., & Beerenwinkel, N. (2018). Network-based integration of multi-omics data for prioritizing cancer genes. Bioinformatics, 34(14), 2441–2448. https://doi.org/10.1093/bioinformatics/bty148

Fujita, K. A., Ostaszewski, M., Matsuoka, Y., Ghosh, S., Glaab, E., Trefois, C., Crespo, I., Perumal, T. M., Jurkowski, W., Antony, P. M. A., Diederich, N., Buttini, M., Kodama, A., Satagopam, V. P., Eifes, S., del Sol, A., Schneider, R., Kitano, H., & Balling, R. (2013). Integrating pathways of parkinson’s disease in a molecular interaction map. Molecular Neurobiology, 49(1), 88–102. https://doi.org/10.1007/s12035-013-8489-4

Gaulton, A., Hersey, A., Nowotka, M., Bento, A. P., Chambers, J., Mendez, D., Mutowo, P., Atkinson, F., Bellis, L. J., Cibrián-Uhalte, E., Davies, M., Dedman, N., Karlsson, A., Magariños, M. P., Overington, J. P., Papadatos, G., Smit, I., & Leach, A. R. (2016). The ChEMBL database in 2017. Nucleic Acids Research, 45(D1), D945–D954. https://doi.org/10.1093/nar/gkw1074

Girvan, M., & Newman, M. E. J. (2002). Community structure in social and biological networks. Proceedings of the National Academy of Sciences, 99(12), 7821–7826. https://doi.org/10.1073/pnas.122653799

Hasin, Y., Seldin, M., & Lusis, A. (2017). Multi-omics approaches to disease. Genome Biology, 18(1). https://doi.org/10.1186/s13059-017-1215-1

Hoang, Q. Q. (2014). Pathway for Parkinson disease. Proceedings of the National Academy of Sciences, 111(7), 2402–2403. https://doi.org/10.1073/pnas.1324284111

Kinnings, S. L., Liu, N., Tonge, P. J., Jackson, R. M., Xie, L., & Bourne, P. E. (2011). A machine learning-based method to improve docking scoring functions and its application to drug repurposing. Journal of Chemical Information and Modeling, 51(2), 408–419. https://doi.org/10.1021/ci100369f

Kuang, Z., Bao, Y., Thomson, J., Caldwell, M., Peissig, P., Stewart, R., Willett, R., & Page, D. (2018). A machine-learning-based drug repurposing approach using baseline regularization. In Methods in Molecular Biology (pp. 255–267). Springer New York. http://dx.doi.org/10.1007/978-1-4939-8955-3_15

Kushwaha, S. K., & Shakya, M. (2010). Protein interaction network analysis—Approach for potential drug target identification in Mycobacterium tuberculosis. Journal of Theoretical Biology, 262(2), 284–294. https://doi.org/10.1016/j.jtbi.2009.09.029

Lee, A., & Gilbert, R. M. (2016). Epidemiology of parkinson disease. Neurologic Clinics, 34(4), 955–965. https://doi.org/10.1016/j.ncl.2016.06.012

Lee, T. K., & Yankee, E. L. (2022). A review on Parkinson’s disease treatment. Neuroimmunology and Neuroinflammation, 8, 222. https://doi.org/10.20517/2347-8659.2020.58

Moriwaki, H., Tian, Y.-S., Kawashita, N., & Takagi, T. (2018). Mordred: A molecular descriptor calculator. Journal of Cheminformatics, 10(1). https://doi.org/10.1186/s13321-018-0258-y

Mysinger, M. M., Carchia, M., Irwin, John. J., & Shoichet, B. K. (2012). Directory of useful decoys, enhanced (DUD-E): Better ligands and decoys for better benchmarking. Journal of Medicinal Chemistry, 55(14), 6582–6594. https://doi.org/10.1021/jm300687e

Patrick, M. T., Raja, K., Miller, K., Sotzen, J., Gudjonsson, J. E., Elder, J. T., & Tsoi, L. C. (2019). Drug Repurposing Prediction for Immune-Mediated Cutaneous Diseases using a Word-Embedding–Based Machine Learning Approach. Journal of Investigative Dermatology, 139(3), 683–691. https://doi.org/10.1016/j.jid.2018.09.018

Pedregosa, F., Varoquaux, G., Gramfort, A., Michel, V., Thirion, B., Grisel, O., Blondel, M., Müller, A., Nothman, J., Louppe, G., Prettenhofer, P., Weiss, R., Dubourg, V., Vanderplas, J., Passos, A., Cournapeau, D., Brucher, M., Perrot, M., & Duchesnay, É. (2012, January 2). Scikit-learn: Machine learning in python. ArXiv.Org. https://arxiv.org/abs/1201.0490

Peng, Q., & Schork, N. J. (2014). Utility of network integrity methods in therapeutic target identification. Frontiers in Genetics, 5. https://doi.org/10.3389/fgene.2014.00012

Poewe, W., Seppi, K., Tanner, C. M., Halliday, G. M., Brundin, P., Volkmann, J., Schrag, A.-E., & Lang, A. E. (2017). Parkinson disease. Nature Reviews Disease Primers, 3(1). https://doi.org/10.1038/nrdp.2017.13

Saito, R., Smoot, M. E., Ono, K., Ruscheinski, J., Wang, P.-L., Lotia, S., Pico, A. R., Bader, G. D., & Ideker, T. (2012). A travel guide to Cytoscape plugins. Nature Methods, 9(11), 1069–1076. https://doi.org/10.1038/nmeth.2212

Shannon, P., Markiel, A., Ozier, O., Baliga, N. S., Wang, J. T., Ramage, D., Amin, N., Schwikowski, B., & Ideker, T. (2003). Cytoscape: A software environment for integrated models of biomolecular interaction networks. Genome Research, 13(11), 2498–2504. https://doi.org/10.1101/gr.1239303

Song, B., Sridhar, P., Kahveci, T., & Ranka, S. (2009). Double iterative optimisation for metabolic network-based drug target identification. International Journal of Data Mining and Bioinformatics, 3(2), 124. https://doi.org/10.1504/ijdmb.2009.024847

Sridhar, P., Kahveci, T., & Ranka, S. (2006, December). AN ITERATIVE ALGORITHM FOR METABOLIC NETWORK-BASED DRUG TARGET IDENTIFICATION. Biocomputing 2007. http://dx.doi.org/10.1142/9789812772435_0009

Subramanian, I., Verma, S., Kumar, S., Jere, A., & Anamika, K. (2020). Multi-omics data integration, interpretation, and its application. Bioinformatics and Biology Insights, 14, 117793221989905. https://doi.org/10.1177/1177932219899051

Surmeier, D. J., Schumacker, P. T., Guzman, J. D., Ilijic, E., Yang, B., & Zampese, E. (2017). Calcium and Parkinson’s disease. Biochemical and Biophysical Research Communications, 483(4), 1013–1019. https://doi.org/10.1016/j.bbrc.2016.08.168

Szklarczyk, D., Gable, A. L., Nastou, K. C., Lyon, D., Kirsch, R., Pyysalo, S., Doncheva, N. T., Legeay, M., Fang, T., Bork, P., Jensen, L. J., & von Mering, C. (2020). The STRING database in 2021: Customizable protein–protein networks, and functional characterization of user-uploaded gene/measurement sets. Nucleic Acids Research, 49(D1), D605–D612. https://doi.org/10.1093/nar/gkaa1074

Tanoli, Z., Vähä-Koskela, M., & Aittokallio, T. (2021). Artificial intelligence, machine learning, and drug repurposing in cancer. Expert Opinion on Drug Discovery, 16(9), 977–989. https://doi.org/10.1080/17460441.2021.1883585

Toodayan, N. (2018). James Parkinson’s Essay on the shaking palsy, 1817–2017. Medical Journal of Australia, 208(9), 384–386. https://doi.org/10.5694/mja17.01085

Vidyadhara, D. J., Lee, J. E., & Chandra, S. S. (2019). Role of the endolysosomal system in Parkinson’s disease. Journal of Neurochemistry, 150(5), 487–506. https://doi.org/10.1111/jnc.14820

Wu, G., Dawson, E., Duong, A., Haw, R., & Stein, L. (2014). ReactomeFIViz: A Cytoscape app for pathway and network-based data analysis. F1000Research, 3, 146. https://doi.org/10.12688/f1000research.4431.2

Wu, L., Shen, Y., Li, M., & Wu, F.-X. (2015). Network output controllability-based method for drug target identification. IEEE Transactions on NanoBioscience, 14(2), 184–191. https://doi.org/10.1109/tnb.2015.2391175

Xing, H., & Gardner, T. S. (2006). The mode-of-action by network identification (MNI) algorithm: A network biology approach for molecular target identification. Nature Protocols, 1(6), 2551–2554. https://doi.org/10.1038/nprot.2006.300

